# Distinct microbial communities alter litter decomposition rates in a fertilized coastal plain wetland

**DOI:** 10.1101/732883

**Authors:** Megan E. Koceja, Regina B. Bledsoe, Carol Goodwillie, Ariane L. Peralta

**Author notes:** Author contributions: MEK, RBB, CG, and ALP conceived and designed the research; MEK, RBB, and ALP collected and analyzed the data; MEK and ALP wrote the manuscript; all authors performed field work and edited the manuscript.

## Abstract

Human activities have led to increased deposition of nitrogen (N) and phosphorus (P) into soils. Nutrient enrichment of soils is known to increase plant biomass and rates of microbial litter decomposition. However, interacting effects of hydrologic position and associated changes to soil moisture can constrain microbial activity and lead to unexpected nutrient feedbacks on microbial community structure-function relationships. Examining how feedbacks of nutrient enrichment on decomposition rates is essential for predicting microbial contributions to carbon (C) cycling as atmospheric deposition of nutrients persists. This study explores how long-term nutrient addition and contrasting litter chemical quality influence soil bacterial community structure and function. We hypothesize that long-term nutrient enrichment of low fertility soils alters bacterial community structure and leads to higher rates of litter decomposition with decreasing C:N ratio of litter; but low nutrient and dry conditions limit constrain microbial decomposition of high C:N ratio litter. We leverage a long-term fertilization experiment to test how nutrient enrichment and hydrologic manipulation (due to ditches) affects decomposition and soil bacterial community structure in a nutrient poor coastal plain wetland. We conducted a litter bag experiment and characterized litter-associated and bulk soil microbiomes using 16S rRNA bacterial sequencing and quantified litter mass losses and soil physicochemical properties. Results revealed that distinct bacterial communities were involved in decomposing higher C:N ratio litter more quickly in fertilized compared to unfertilized especially under drier soil conditions, while decomposition rates of green tea litter (lower C:N ratio) were similar between fertilized and unfertilized plots. Bacterial community structure in part explained litter decomposition rates, and long-term fertilization and drier hydrologic status affected bacterial diversity and increased decomposition rates. However, community composition associated with high C:N litter was similar in wetter plots with available nitrate detected, regardless of fertilization treatment. This study provides insight into long-term fertilization effects on soil bacterial diversity and composition, decomposition, and the increased potential for soil C loss as nutrient enrichment and hydrology interact to affect historically low nutrient ecosystems.

## Introduction

Humans modify their landscapes through fossil fuel burning, deforestation, and intense agricultural activity (Vitousek et al. 2010, Fowler et al. 2013). These anthropogenic disturbances have led to increased atmospheric deposition of nitrogen (N) and phosphorus (P), and can be particularly disruptive to historically nutrient-limited ecosystems (Guignard et al. 2017). This increased nutrient deposition can cause a fertilization effect on plant-microbial interactions which results in increased biomass and shifts in community structure (Cherif and Loreau 2009, Leff et al. 2015, Harpole et al. 2016). This fertilization effect can increase plant biomass C, which fuels heterotrophic microbial growth and leads to increased respiration of CO_2_ (Hoosbeek et al. 2004, Kuzyakov 2010). The extent to which nutrient enrichment predicts decomposition rates is also determined by the interaction of soil microorganisms with the plant inputs and the abiotic soil environment. While human alteration of the environment and its effect on C storage and release is relatively well-documented in terrestrial ecosystems, the mechanisms governing the interactive effects of nutrient enrichment on plant-soil-microbial relationships at terrestrial-aquatic interfaces are relatively understudied. The availability of terminal electron acceptors in hydrologically dynamic ecosystems are known to shape the resident microbial community and to control biogeochemical functions (Bernhardt et al. 2017). This makes it challenging to predict microbial responses to environmental change in ecosystem models.

Changes in the nutrient stoichiometry of both plants and surrounding soils are expected to affect microbial community composition and function, leading to changes in elemental cycling. Prior studies have shown that long-term nutrient addition enhances grassland plant biomass and increases C inputs into soils (e.g., Harpole et al. 2016). These additional plant inputs also increase CO_2_ outputs via microbial respiration (Hoosbeek et al. 2004, Lange et al. 2015). If microbes are not limited by N and P, the increase in plant-derived organic C inputs are more quickly metabolized and soil CO_2_ fluxes increase (Peralta and Wander 2008, Cotrufo et al. 2013, Castellano et al. 2015). Plant biomass could also vary in C:N or C:P ratios, which could also cause changes in microbial nutrient use and respiration rates. Thus, it is important to understand how changes in available nutrients within plant litter alter microbial communities and CO_2_ respiration rates as surrounding environmental conditions also shift (Hoosbeek et al. 2004, Lange et al. 2015).

Nuanced interactions between litter chemical quality, abiotic environment, and resident microbial composition can manifest in similar decomposition patterns (i.e., increased rates due to increased nutrient availability). However, the degree to which microbial structure contributes to the magnitude of litter mass loss is challenging to pinpoint since interactions among abiotic (temperature, nutrient availability, moisture) and biotic (community structure) factors determine decomposition rates (Kuzyakov and Blagodatskaya 2015, Meier et al. 2017, Deveau et al. 2018). In this study, we examine the extent that litter-associated microbiomes are unique depending on historical nutrient enrichment and how this community composition contributes to decomposition of litter of varying chemical quality (i.e., C:N ratio).

The nutrient availability in upland conditions makes priming (using standing litter nutrients) more pronounced as aerobic respiration is more energetically favored. In this study, we measured the extent that microbial community change (due to nutrient enrichment) contributes to enhanced decomposition using a model litter comparison. The Tea Bag Index (TBI) compares the decomposition rates of two different plant litters, green and rooibos Lipton™ tea bags (Keuskamp et al. 2013). Green tea has a measured mean C:N ratio of 12.229 while rooibos tea is measured to have a much higher mean C:N ratio of 42.870. We used the TBI protocol to examine how long-term fertilization affects decomposition rates in a nutrient-limited coastal plain wetland ecosystem. We also characterized the bacterial communities (using targeted amplicon sequencing of the V4 region of the 16S rRNA gene) associated with the litter decomposition of two differing litter types (green and rooibos teas) occurring in wetland soils exposed to fertilization or not. The extent that community composition matters to decomposition rate depends on biotic and abiotic factors. In this study, litter type, nutrient enrichment and hydrologic status resulted in distinct bacterial communities.

## Materials and Methods

### Experimental design of a long-term ecological experiment

Initiated in 2003, a long-term ecological experiment started at East Carolina University’s West Research Campus (WRC) (35.6298N, −77.4836W) examines the effects of mowing and fertilization on coastal plain wetland plant and microbial communities. This study site is classified as a jurisdictional wetland, and the plant community has been described as a mosaic of wet pine flatwood habitat, pine savanna, and hardwood communities (Chester 2004). The soils were characterized as fine, kaolinitic, thermic Typic Paleaquults (Coxville series), fine-loamy, siliceous, semiactive, thermic Aeric Paleaquults (Lynchburg series), and fine-loamy, siliceous, subactive, thermic Aquic Paleudults (Goldsboro series). These soils are moderately to poorly drained ultisols (USDA NRCS 2019). The annual mean temperature is 16.4 °C and annual precipitation is 126 cm (U.S. Climate Data 2019). Fertilization and mowing treatments are replicated on eight 20×30 m blocks in a full factorial design, and the N-P-K 10-10-10 pellet fertilizer is applied 3× per year (February, June, and October) for a total annual supplementation of 45.4 kg/ha for each nutrient. Plots are mowed by bush-hog annually to simulate fire disturbance (Goodwillie et al. In Review). Within each plot, annual soil and plant sampling takes place at three randomly-placed quadrats (Fig. 1). The plant community in plots that are mowed is dominated by perennial forbs in two major genera of the Asteraceae—*Eupatorium* and *Solidago*—and graminoid species. The relative abundance of major grass species differs between mowed and fertilized plots, with switchcane (*Arundinaria tecta*) most dominant in fertilized plots and broomsedge (*Andropogon virginicus*) found almost exclusively in unfertilized plots. Sedges (e.g., *Rhychospora* and *Carex* spp.) and species of *Juncus* are common in (wet) plots away from, but not in (dry) plots adjacent to the drainage ditch. This hydrologic gradient is caused by a roadside drainage ditch such that four blocks near the ditch are drier and four blocks away from the ditch are wetter (Goodwillie et al. In Review). In 2018, volumetric soil moisture content (measured by capacitance), was >2 times wetter in blocks away from the ditch compared to blocks adjacent to the ditch. However, this hydrologic gradient has not yet been characterized by modeling flow according to water levels over time. The ditch was not intentionally manipulated as part of our experimental design, but it contributes an important ecological variable to the study (Goodwillie et al. In Review). For this study, tea litter decomposition experiments were focused in the mowed plots only.

**Figure 1.**
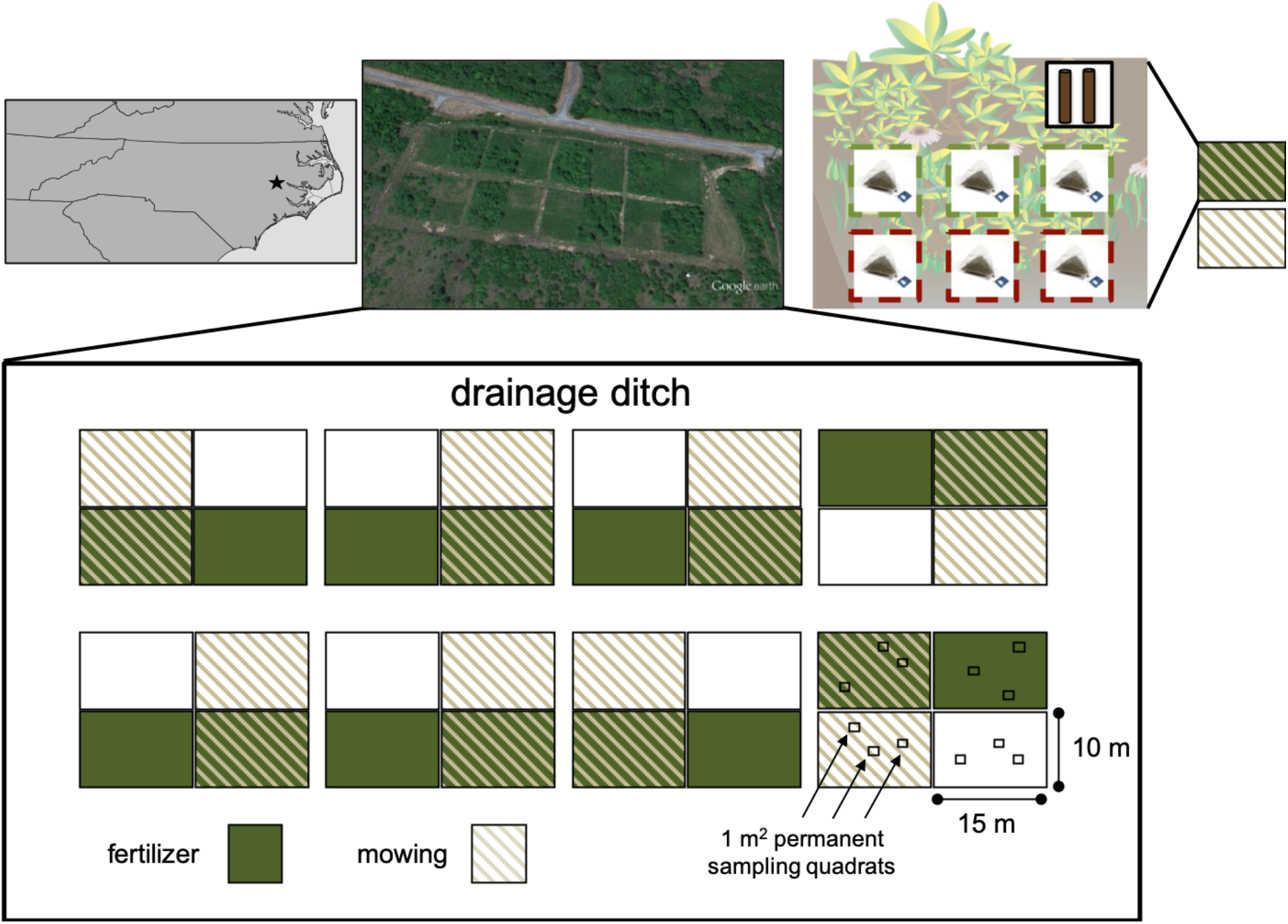
Experimental design of a long-term ecological experiment to test the effects of fertilizer and disturbance by mowing on plant and microbial communities at East Carolina University’s West Research Campus (WRC), Greenville, NC, USA. The decomposition protocol for this experiment was adapted from the Tea Bag Index Protocol (Keuskamp et al. 2013)(Keuskamp et al., 2013) and applied within the eight-replicate fertilized and unfertilized plots of the WRC. In one quadrat per replicate plot, three bags of Lipton™ green tea and three bags of Lipton™ rooibos tea were buried and retrieved after 111 days. Bulk soil was sample as a composite sample representing two soil cores from three permanent quadrats per plot.

### Soil sampling

We collected composite soil samples from all mowed/unfertilized and mowed/fertilized plots, which represented two soil cores (12 cm depth, 3.1 cm diameter) adjacent to each of three permanently installed 1 m^2^ quadrats where plant community data are annually collected. Each composite bulk soil sample was passed through a 4 mm sieve and homogenized before further analysis. This bulk soil sampling occurred on November 14, 2019, about three months after the tea litter bags were collected from the field.

### Soil physico-chemical analyses

We measured gravimetric soil moisture by weighing 20-30 g of field-moist soil, drying at 105 °C overnight, and then re-weighing. We calculated percent moisture as difference between field-moist and dried soils divided by the oven-dried soil weight. In addition, we measured pH of oven-dried soil by mixing a 1:1 soil:water solution and using a pH probe (Genemate-Bioexpress; Kaysville, Utah; USA). We measured total soil organic carbon and total nitrogen on finely ground dried soil using an elemental analyzer (2400 CHNS Analyzer; Perkin Elmer; Waltham, Massachusetts, USA) at the Environmental and Agricultural Testing Service laboratory (Department of Crop and Soil Sciences at North Carolina State University). Extractable inorganic N for each soil sample was measured on approximately 5 g of field moist soil. We added 45 ml of 2 M KCl to soil, shook the sample for about 1 hour, and gravity filtered. Total phosphate (PO_4_^3−^) was extracted by combining 0.1 g dried soil (ground and passed through a 500 μm sieve) with 0.5 ml of 50% w/v Mg(NO_3_) and ashing for 2 hours at 550 °C. Samples were hydrated with 10 mL of 1 M HCl, shaken for 16 hours at 250 RPM, and the filtered (22 μm filter). Water extractable PO_4_^3−^ was determined by combining 1 g dried soil (ground and passed through a 500 μm sieve) with deionized water, shaken for 1 hour, and filtered (22 μm filter). Ammonium (NH_4_^+^), nitrate (NO_3_^−^), and PO_4_^3−^ ions in soil extracts were colorimetrically measured using a SmartChem 200 auto analyzer (Unity Scientific Milford, Massachusetts, USA) at the East Carolina University Environmental Research Laboratory.

### Field experimental methods

The field protocol for this experiment was adapted from the Tea Bag Index Protocol (Keuskamp et al. 2013) and applied within the eight-replicate fertilized and unfertilized plots of the WRC. The experimental setup involved three bags of Lipton™ green tea and three bags of Lipton™ rooibos tea per replicate plot (3 bags × 8 blocks × 2 treatment plots = 48 bags per each tea bag type). In each treatment plot, we prepared six holes about 20 cm apart and 8 cm deep using a hand trowel. We weighed green and rooibos tea bags and buried them in separate holes. Soil was lightly packed around the tea bags, while keeping the labels visible above the soil. Tea bags were recovered after 111 days (May 21–September 09, 2018) using hand trowels to loosen soil adjacent to tea bag locations. Two of three green tea bags at each replicate plot were measured for mass loss. The third green tea bag was placed into a sterile Whirl-pak bag and stored at −20 °C for tea-associated bacterial community analysis. The rooibos tea was processed following the same procedure. Upon returning to the lab, the tea bags for decomposition analysis were separated from adhered soil particles, placed into individual aluminum weighing dishes, and oven-dried at 70 °C for 48 hours. To quantify mass loss, the tea bags were reweighed following drying and compared to their initial weight prior to soil incubation.

### Soil and tea-associated bacterial 16S rRNA sequencing

Following freezing, tea litter was removed from tea bags, and DNA was extracted from samples using the Qiagen DNeasy PowerSoil kit. We also extracted DNA from soils using the Qiagen DNeasy PowerSoil Kit. For each unfertilized or fertilized treatment, DNA was extracted from green tea (15 total), rooibos tea (14 total), and bulk soil samples (16 total). Three litter samples could not be located and were not retrieved from the field. Following extraction, sample DNA was used as template in PCR reactions with a bacterial 515F/806RB barcoded primer set originally developed by the Earth Microbiome Project to target the V4-V5 region of the bacterial 16S subunit of the rRNA gene (Apprill et al., 2015; Caporaso et al., 2012; Parada et al., 2016). For each sample, three 50 μL PCR libraries were prepared by combining 35.75 μL molecular grade water, 5 μL Amplitaq Gold 360 10x buffer, 5 μL MgCl_2_ (25 mM), 1 μL dNTPs (40mM total, 10mM each), 0.25 μL Amplitaq Gold polymerase, 1 μL 515 forward barcoded primer (10 μM), 1 μL 806 reverse primer (10 μM), and 1 μL DNA template (10 ng/μL). Thermocycler conditions for PCR reactions were as follows: initial denaturation (94 °C, 3 minutes); 30 cycles of 94 °C for 45 seconds, 50 °C for 30 seconds, 72 °C for 90 seconds; final elongation (72 °C, 10 minutes). Triplicate PCR reactions were combined for each sample and cleaned according to the Axygen^®^ AxyPrep Magnetic Bead Purification Kits protocol (Corning Life Sciences). Following cleaning, PCR product were quantified using Quant-iT dsDNA BR (broad-range) assay (Thermo Scientific, Waltham, MA, USA). Libraries were pooled in equimolar concentration of 5 ng/μL after being diluted to a concentration of 10 ng/μL. The Indiana University Center for Genomics and Bioinformatics sequenced the pooled libraries using the Illumina MiSeq platform using paired-end reads (Illumina Reagent Kit v2, 500 reaction kit).

Following sequencing, we processed raw sequences using a standard mothur pipeline (v1.40.1) (Schloss et al. 2009, Kozich et al. 2013). Contigs were assembled from paired end reads and quality trimmed using a moving average quality score (minimum score of 35 bp). Sequences were aligned to the SILVA rRNA gene database (v.128) (Quast et al. 2013) and chimeric sequences were removed using the VSEARCH algorithm (Rognes et al. 2016). Formation of operational taxonomic units (OTUs) involved dividing sequences based on taxonomic class and then binnned into OTUs with a 97% sequence similarity level. OTUs were classified using the SILVA database (Yilmaz et al. 2014, Glöckner et al. 2017).

### Statistical analyses

All statistical calculations were completed in the R environment (R v3.6.3, R Core Development Team 2020). Decomposition was measured through mass loss of buried tea bags. Initial and final tea bag weights were used to determine percent of loss. We ran a linear mixed effects model with ‘source’, ‘treatment’, and ‘ditch proximity’ as fixed effects and ‘block’ as a random effect using the lmer() function in the lmerTest package (Kuznetsova et al. 2017). For the mass loss response variable, a linear mixed model was fit by REML and produced type II analysis of variances tables (ANOVA) tables based on the Kenward-Roger’s denominator degrees of freedom method using the lmerModLmerTest() function. Calculations of decomposition rate (*k*) and stabilization factor (*S*) for each tea bag were accomplished according to formulas found within the TBI protocol. Decomposition rate, *k*, is a parameter representing short-term dynamics of new input decomposition, while stabilization factor, *S*, is a parameter characterizing long-term carbon storage. These values were used to draw comparisons between *k* and *S* values found within the WRC wetland ecosystem to those of global ecosystems submitted to the Tea Bag Index protocol (Keuskamp et al. 2013). Outcomes of these calculations were plotted on a graph, and locations of groupings were compared to those of the larger multi-ecosystem index (Keuskamp et al. 2013).

For the bulk soil factors measured and the computed diversity (OTU richness, Shannon Diversity, Simpson’s Evenness) metrics for soil and tea litter associated bacterial communities, we ran a series of linear mixed models. We ran linear mixed effects models with ‘source’, ‘treatment’ and ‘ditch’ as fixed effects and ‘block’ as a random effect and fitted by the REML criterion using the lmer() function in the lmerTest package (Kuznetsova et al. 2017). Statistical inferences for fixed effects were calculated from type II ANOVA tables and the Kenward-Roger’s degrees of freedom method using the anova() function.

To visualize the community responses to fertilization and litter type, we used principal coordinates analysis (PCoA) of bacterial community composition based on the Bray-Curtis dissimilarity. We used a permutational multivariate analysis of variance (PERMANOVA) to examine among-treatment differences in bacterial communities. We also include a comparison of the bulk soil and tea litter associated bacterial communities (PERMANOVA, PCoA ordination) but focused on the specific tea litter associated microbiome patterns. To identify which bacterial species were most representative of each litter type, we ran an indicator species analysis. For the indicator species analysis, we only included bacterial taxa with a relative abundance greater than 0.05 when summed across all plots. We performed PERMANOVA using the adonis() function in the vegan package (Oksanen et al. 2017) and the indval() function in the indicspecies package (Caceres and Jansen 2016).

To examine structure-function relationships between the tea-associated bacterial community and the mass loss of the tea litter, we ran a linear regression to measure the relationship between mass loss as a function of bacterial diversity (mass loss ~ OTU richness + Shannon Diversity + Simpson’s Evenness). In addition, we conducted a distance-based partial least squares regression (dbplsr) measure how much variation in mass loss of litter (function) was explained by the tea-associated bacterial community composition (structure) (mass loss ~ bacterial community). We used the Bray-Curtis dissimilarity matrix and the generalized cross-validation estimate of the prediction error (‘gcv’ method). We performed a distance-based partial least squares regression analysis using the dbplsr() function in the dbstats and pls packages (Boj Del Val et al. 2007, Boj et al. 2017, Mevik et al. 2019).

## Results

Long-term fertilization strongly influenced abiotic and biotic factors within this nutrient-limited wetland environment. Both fertilization and ditch effects influenced soil pH, soil C:N ratio, extractable nitrate concentrations, and water extractable phosphorus (Table S1A, Table S2B). A subset of soil factors was distinct between fertilized and unfertilized soils in the mowed plots. Total and water extractable phosphorus concentrations, extractable nitrate concentrations, soil C and N content, moisture, and pH were higher in fertilized compared to unfertilized soils (Table S1B). In addition, soil moisture, C:N ratio, and water extractable phosphorus were higher in wetter soils away from the ditch compared to drier soils adjacent to the ditch (Table S1C). Finally, nitrate concentrations were detectable only in wetter soils away from the ditch with higher concentrations in fertilized soils compared to unfertilized soils while nitrate was below detection limits in all soils near the ditch (Table S1A). These differences in fertilized compared to unfertilized soils and soil moisture due to ditch proximity related to litter decomposition rates and altered bacterial community structure.

To examine how litter type and nutrient addition influenced decomposition rates, we measured mass loss of green (low C:N litter) and rooibos (high C:N litter) tea bags. Following an 111-day incubation, there was a significant litter type (source) × fertilization treatment interaction (*P*=0.05) (Fig. 2, Table S2). The mass loss of green tea litter was ~24% higher than the overall mass loss of rooibos tea litter (Fig. 2, Table S2), averaged across fertilized and unfertilized soils. Both fertilization and proximity to drainage ditch also increased the rate of mass loss (Fig. 2, Table S2). Across all tea types and drainage ditch proximities, fertilized soils showed an average mass loss of ~7.5% more than that of unfertilized soils. Tea bags buried in closer proximity to the drainage ditch had ~6% more mass loss averaged across all litter and treatment types.

**Figure 2.**
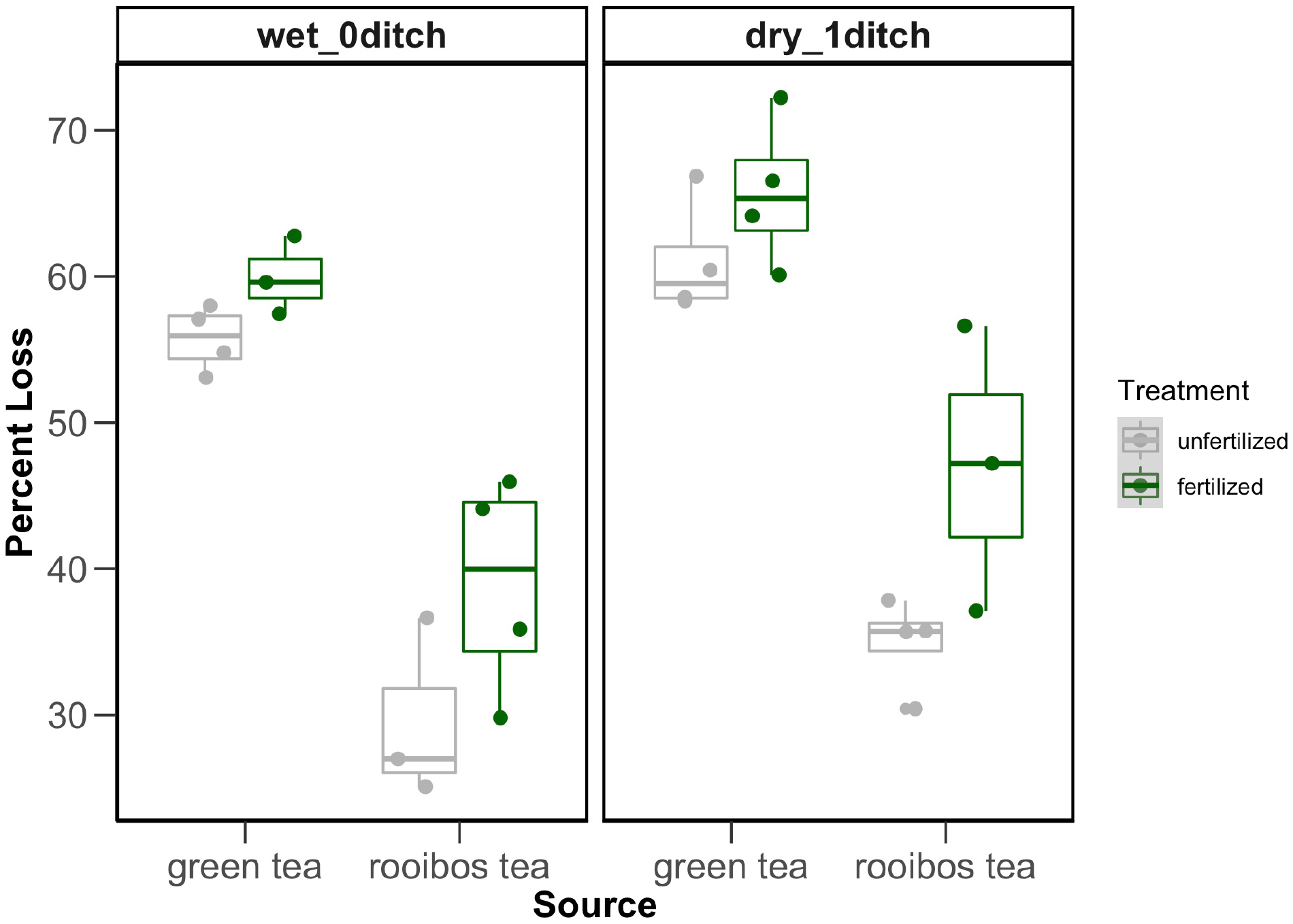
Boxplots representing cumulative mass loss of green (low C:N ratio litter) and rooibos (high C:N ratio litter) tea in fertilized (green) and unfertilized (gray) plots. Symbols represent individual data points.

We also compared the decomposition rates (*k*) and stabilization factors (*S*) from the long-term fertilization experiment to a broader context of ecosystems (Keuskamp et al., 2013). The most variable *S* and *k* values were measured in the wetter, fertilized plots (Fig. 3). The unfertilized compared to fertilized samples were generally higher along the stabilization factor (*S*) axis. Samples that were buried in the wetter, fertilized plots clustered along the higher end of the stabilization factor (*S*), while litter buried in the drier plots (fertilized and unfertilized) had lower estimated stabilization factor (*S*) (Fig. 3). Upon comparison with other ecosystems, a large number of fertilized treatment samples grouped near the grassland-ambient ecosystem in Iceland and the forest ecosystem in the Netherlands (Keuskamp et al. 2013). However, the unfertilized treatment samples grouped near the peat-disturbed and peat-undisturbed ecosystems of Iceland and the peat ecosystem of the Netherlands (Keuskamp et al. 2013). Within this coastal plain wetland system, long-term nutrient enrichment shifted decomposition rates that were representative of grassland and forested ecosystems (fertilized plots) or peatlands (unfertilized plots) (Fig. 3).

**Figure 3.**
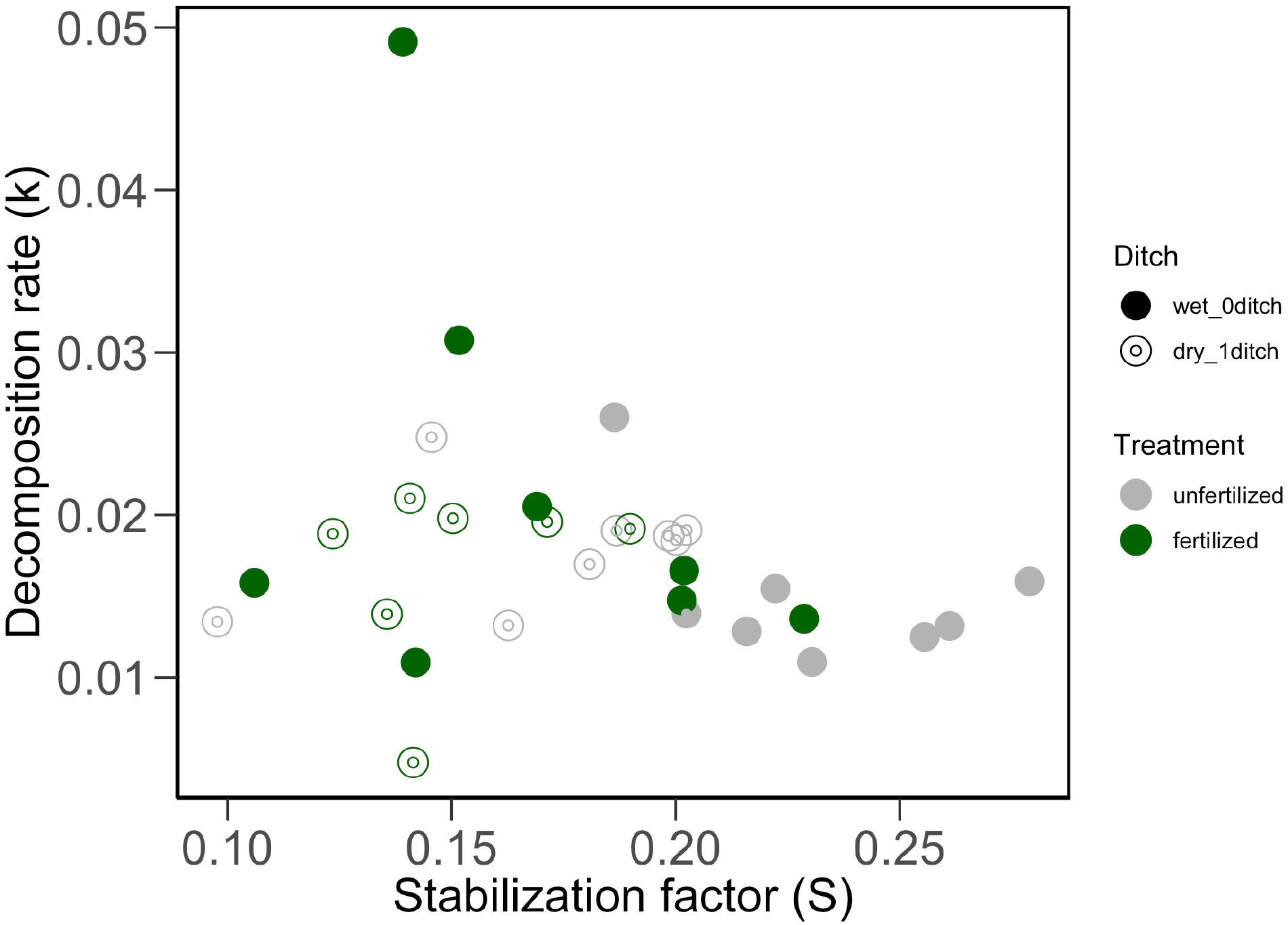
Initial decomposition rate *k* and stabilization factor *S* for different tea bags within the long-term fertilization experiment. Tea bags from unfertilized plots are indicated in grey, while tea bags from fertilized plots are indicated in green. Open symbols represented drier plots that were closer to the drainage ditch, while closed symbols represented wetter plots that were further from the ditch.

Fertilization effects on soil physicochemical properties and decomposition rates were related to shifts in bulk soil and litter-associated bacterial communities. Fertilization effect, tea type, and proximity to ditch (i.e., wetter vs. drier plots) influenced bacterial diversity in various ways. Specifically, tea type (*R*^*2*^=0.130, *P*=0.001) influenced bacterial community composition the most, while the interaction of tea type and proximity to ditch affected bacterial community patterns to a lesser degree (*R*^*2*^=0.060, =0.002) (Fig. 4, Table S3) (see circle vs triangle). The proximity to ditch (proxy for hydrology) also altered bacterial community composition (*R*^*2*^=0.110, *P*=0.001) (Fig. 4, Table S3) (see opened vs filled). Within tea type, fertilization treatment influenced bacterial community composition (*R*^*2*^=0.070, *P*=0.003) (Fig. 4, Table S3) (see green vs gray). For the drier, ditched plots, bacterial communities associated with high C:N ratio litter (rooibos) overlapped despite fertilization treatment, while bacterial communities were distinct between fertilization treatments in the wetter plots (Fig. 4). When the bulk soil and tea-associated bacterial communities were analyzed together, bulk soil or tea most strongly influenced bacterial community composition (source: *R*^*2*^=0.490, *P*=0.001) (Fig. S1, Table S4). In addition to these observed patterns, OTU diversity was generally higher in fertilized compared to unfertilized plots, especially in the drier plots associated with the drainage ditch (Fig. 5, Table S5). Specifically, OTU richness values were highest in bulk soil compared to tea types (Fig. 5A, Table S5A), and Shannon diversity values were highest in bulk soil compared to tea type in fertilized plots (Fig. 5B, Table S5B). Lastly, bacterial evenness was similar across fertilization treatment, source type, and proximity to ditch (Fig. 5C, Table S5C).

**Figure 4.**
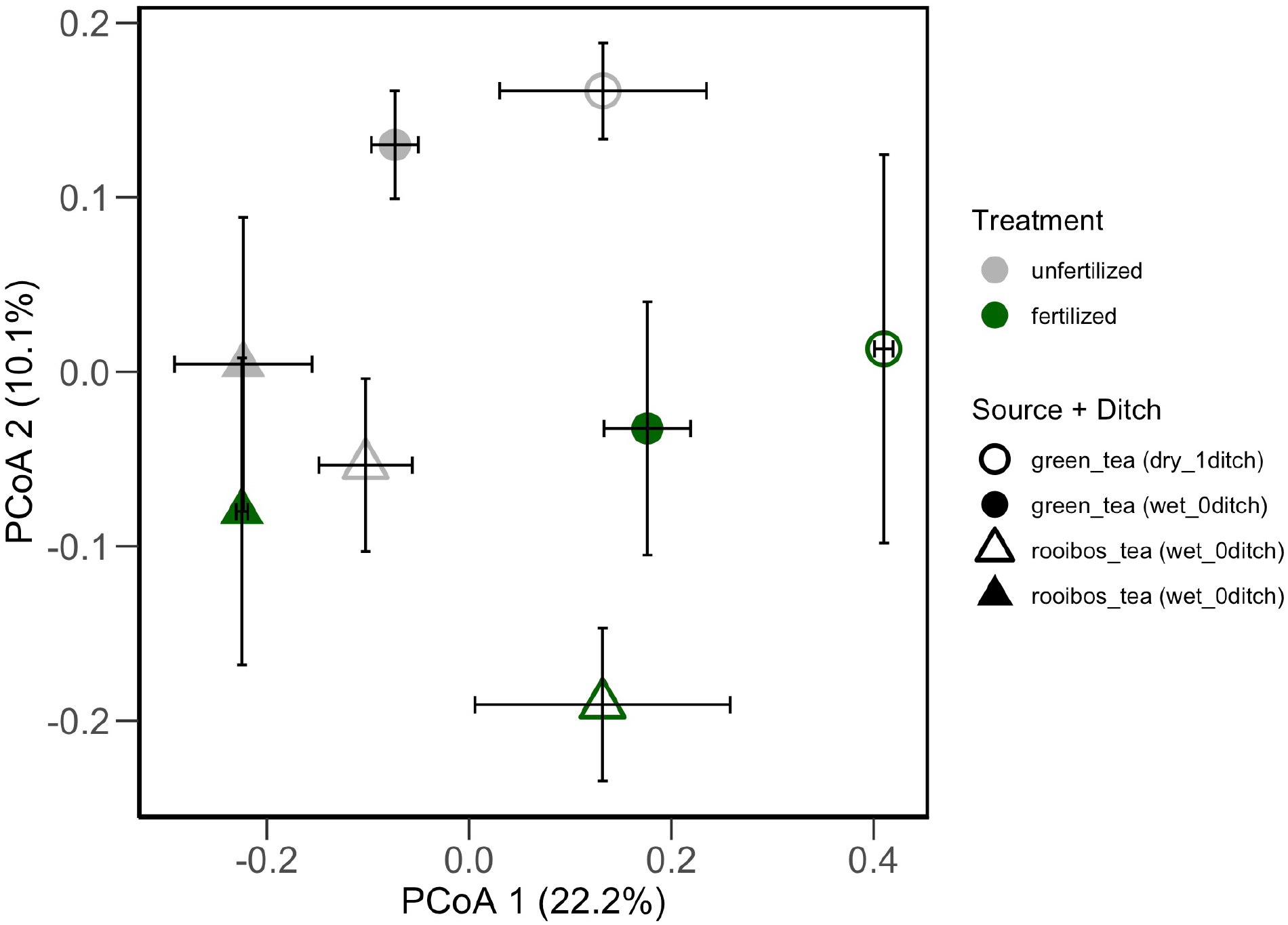
Ordination based on a Principal Coordinates Analysis depicting bacterial community composition according to tea type. Symbols are colored according to fertilization treatment (gray = unfertilized, green = fertilized) and tea source (circles = low C:N ratio green tea, triangles = high C:N ratio rooibos tea) at drier mowed plots situated close to the drainage ditch (open symbols) compared to wetter mowed plots (closed symbols).

**Figure 5.**
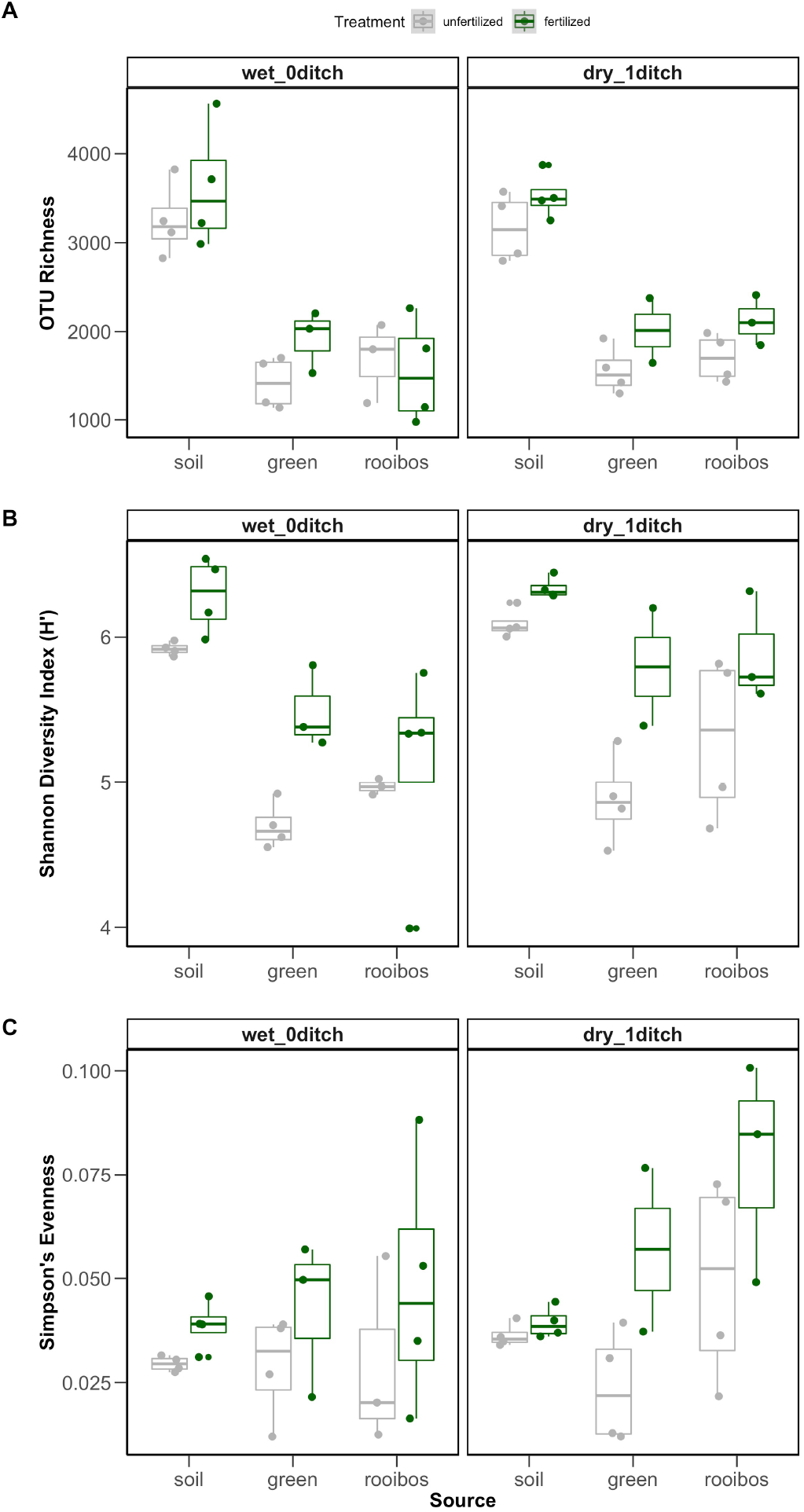
Boxplots depicting of bacterial diversity metrics (OTU richness, Shannon Diversity Index H′, and Simpson’s Evenness) associated with source (bulk soil, low C:N ratio green tea, high C:N ratio rooibos tea). Boxplots and symbols are colored according to fertilization treatment (gray = unfertilized, green = fertilized) at drier mowed plots situated close to the drainage ditch compared to wetter mowed plots. Symbols represent data points for individual plot samples.

To further examine bacterial associations with litter decomposition, we used indicator species analysis to identify a subset of bacterial taxa that represented each of the bulk soil or tea associated microbiomes. The unclassified OTU within the class Spartobacteria and another OTU within the order Acidobacteria Gp1 represented the unfertilized bulk soil bacterial community, while an unclassified OTU within the order Solirubrobacterales represented bulk soils in the fertilized, dry plots. In wetter plots, an unclassified OTU within the order Rhodospirillales represented unfertilized plots while unclassified taxa within the orders Acidobacteria Gp1, Gp2, and Gp6 signified fertilized bulk soils (Tables S6, S7). In addition, the unclassified OTU within the Acidobacteria Gp3 and *Dongia* spp. represented the green tea-associated (low C:N litter) bacterial communities in the unfertilized, dry plots while *Conexibacter* spp. were represented in the fertilized, dry plots. In the wetter plots, *Phenylobacterium* spp. represented green tea-associated bacterial communities in unfertilized plots and *Legionella* spp. in fertilized plots. The OTUs *Acidisoma* spp. in unfertilized plots and *Dyella* spp. in fertilized characterized rooibos tea associated bacterial communities in the dry plots. Lastly, the OTU *Lacibacterium* spp. represented mowed plots, while *Dokdonella* spp. and an unclassified OTU within the genus *Microbacteriaceae* represented rooibos tea associated communities in the wet plots (Tables S6, S7).

When examined together, the link between bacterial community structure and decomposition function was relatively weak. No relationship between litter mass loss and bacterial diversity was detected (*R*^*2*^_*adj*_ = −0.038, *P* = 0.574). However, when the total bacterial community was considered, the tea-associated bacterial community accounted for ~64 % of the variation in litter mass loss (dbplsr, Component 1 *R*^*2*^_*adj*_ = 63.56, Component 2 *R*^*2*^_*adj*_ = 87.09, Component 3 *R*^*2*^_*adj*_ = 93.96) (Table 1).

**Table 1.** Summary of distance-based partial least squares regression (dbplsr) representing how much variation in decomposition rate is explained (%) by each component (Comp) derived from a tea-associated bacterial community Bray-Curtis distance matrix.

|  | Comp 1 | Comp 2 | Comp 3 | Comp 4 | Comp 5 | Comp 6 |
| --- | --- | --- | --- | --- | --- | --- |
| $R^2$ | 64.96 | 88.08 | 94.65 | 98.12 | 99.32 | 99.77 |
| adjusted $R^2$ | 63.56 | 87.09 | 93.96 | 97.78 | 99.16 | 99.70 |
| gvar | 31.50 | 41.62 | 50.10 | 56.93 | 61.21 | 66.39 |
| crit | 2.47 | 0.91 | 0.44 | 0.17 | 0.07 | 0.03 |

gvar = total weighted geometric variability; crit = value of criterion defined in method

## Discussion

Litter composition, fertilization, and ditch effects on soil moisture influenced bacterial community structure-function relationships in unexpected ways. In this study, increases in litter mass loss were greater in fertilized compared to unfertilized soils, especially in drier versus wetter plots. Surprisingly, distinct microbial communities associated with different litter chemical qualities. Litter mass loss following the field incubation indicates that similar bacterial communities are capable of decomposing higher C:N ratio litter (rooibos tea) more quickly in fertilized compared to unfertilized plots, particularly in drier compared to wetter plots. However, lower C:N ratio litter (green tea) decomposition rates were similar among fertilized and unfertilized plots, although slightly higher in drier compared to wetter plots but were represented by distinct bacterial communities. As expected, the low C:N ratio litter provides a N source to microbes regardless of external soil nutrient conditions, whereas the high C:N ratio litter is reliant on N from the fertilized soil in order to support increased rates of litter decomposition (Duddigan et al. 2020). These results suggest that shifts in microbial community composition are partly due to differences in litter chemical quality and nitrate availability in soils.

The comparative results from the TBI-based decomposition experiment indicate that the tea litter bags buried in the unfertilized treatment had decomposition rates (*k*) and C stabilization factor (S) values similar to that of peatland ecosystems (Keuskamp et al. 2013). Lower decomposition rates occurred in unfertilized plots (i.e., the ambient state of this nutrient-poor habitat). When looking specifically at high C:N ratio litter, less mass was lost to decomposition. This provides some evidence that C storage potential within these plots resembles that of peatlands (Hill et al. 2018). Wetland ecosystems that are disconnected and isolated from pulses of nutrients due to natural processes or agricultural or urban runoff are more commonly nutrient limited by N and P (Vitousek et al. 2010). The plant species that are adapted to these low-nutrient ecosystems can maintain positive population growth and contribute to organic C additions to soils. Further, flooded environments, such as wetlands, are also known to support anaerobic microbial processes, which result in slower rates of decomposition (Collins et al. 2015). Taken together, low nutrient environments can be sites of high plant biodiversity leading to organic C inputs and balanced with slower decomposition rates, which contribute to long-term C storage in soils (Hooper et al. 2005, Kleber et al. 2011). These results reflect the coastal plain wetland environment in this study, which offers some evidence that wetland environments store a disproportionate amount of C compared to other ecosystems (Sutfin et al. 2016, Nahlik and Fennessy 2016). However, this C storage capacity can be disrupted by nutrient enrichment. The fertilized plots, however, had *k* and *S* values that grouped closer towards grassland and forest ecosystems (Riggs et al. 2015), which have lower C storage potential because more available nutrients and oxic environments support aerobic respiration and higher decomposition rates. Results from this study provide support that nutrient enrichment can have a lasting influence on plant-soil-microbe interactions that affect C storage potential of wetland ecosystems (Lambers et al. 2009, Allison et al. 2014, Hartman et al. 2017).

Patterns in bacterial community composition (i.e., beta diversity) but not the combined OTU richness, Shannon diversity, Simpson’s evenness diversity metrics, significantly explained decomposition rates. Further, litter composition followed by the fertilization and ditch effects determined bacterial community composition. A trend of higher decomposition rates was associated with high bacterial diversity, especially in the fertilized compared to unfertilized plots. Higher diversity has been associated with higher decomposition rates across different ecosystems (Lange et al. 2015, Trivedi et al. 2016). Shifts in plant and microbial communities can fuel decomposition rates and ultimately C losses from historically low nutrient ecosystems that experience atmospheric nutrient deposition (Sardans and Peñuelas 2012, Wieder et al. 2015).

When bulk and tea-associated microbiomes were considered together, the bulk soil bacterial communities were very distinct from the tea-associated bacterial communities. Within the bulk soil, bacterial communities were distinct across fertilized and unfertilized dry plots, and indicator taxa were phylogenetically distinct at the phylum level (Proteobacteria vs. Acidobacteria). While results revealed similarity in bulk soil bacterial composition in the fertilized and unfertilized wet plots, the indicator taxa also represented distinct phyla (unfertilized: Verrucomicrobia + Acidobacteria, fertilized: Actinobacteria). In contrast, the indicator taxa for tea-associated bacterial communities that overlapped in composition were similar at the phylum level. Specifically, similar bacterial assemblages of rooibos tea-associated (high C:N litter) microbiomes in the fertilized and unfertilized, dry plots were associated with Proteobacteria (Table S6, unfertilized: *Lacibacterium spp.*, fertilized: *Dokdonella spp.*). Indicator bacterial taxa representing high C:N ratio litter-associated microbiomes in fertilized, dry plots were *Dokdonella spp.* (Proteobacteria phylum) and unclassified Microbacteriaceae (Actinobacteria phylum). The genus *Dokdonella* has been involved in heterotrophic denitrification (Figueroa-González et al. 2016, Palma et al. 2018), while members of the family Microbacteriaceae have been predominantly found in terrestrial and aquatic ecosystems with putative functions related to plant pathogenicity (Glöckner et al. 2000, Young 2008, Lory 2014). Indicator taxa within the order Alphaproteobacteria represented the similar bacterial communities of the high C:N litter-associated microbiomes in unfertilized (indicator OTU *Acidisoma spp.*), wet plots and low C:N litter-associated microbiomes in unfertilized, dry plots (indicator OTU *Phenylobacterium spp.*). In addition, the community similarity between low C:N litter-associated microbiomes in unfertilized, wet plots and high C:N litter-associated microbiomes in fertilized, wet plots were represented by taxa within different Proteobacterial classes. Specifically, *Dongia spp.* (within Alphaproteobacteria class) and *Dyella spp.* (within Gammaproteobacteria class) are similar in at least some life history traits. Representative isolates were cultured from soils (*Dyella*) and wetland sediments (*Dongia*) and were found to be aerobic, Gram-negative, and motile (Xie and Yokota 2005, Baik et al. 2013). Further, these assemblages also overlapped with green tea-associated microbiomes in the fertilized, dry plots, which were represented by *Legionella spp.* (within class Gammaproteobacteria). Legionellae are found in aquatic and other moist environments with their free-living protozoan hosts (Barker and Brown 1994). In the wetland soil environment, they can persist but are unlikely existing under optimal conditions.

The interaction between source (litter type) and fertilization treatment influenced mass loss of litter but hydrologic setting and soil nitrate altered litter-associated microbiomes to varying degrees. The ditch-derived hydrologic differences at this wetland resulted in different structure-function relationships for high C:N ratio litter-associated microbiomes only. Under the relatively dry plots (i.e., near drainage ditch), microbiomes associated with high C:N ratio litter were similar, but mass loss of litter was higher in fertilized compared to unfertilized plots. In all other instances, distinct, fertilized microbiomes were associated with higher decomposition rates compared to unfertilized plots. In the case that community structure is the same and decomposition rates increase, this pattern provides some evidence of more severe N limitation in drier compared to wetter plots, where soil nitrate concentrations are below detection limits (Table S1B). This result was only observed under high C:N ratio litter context since the low C:N litter provided much needed N to litter-associated microbes. While soil ammonium levels are similar in this case, it is possible that N mineralization is occurring without limitation, but nitrification processes could be slowed due to low soil pH, resulting in differences in extractable soil nitrate but not ammonium concentrations (Hinckley et al. 2019).

In this study, nutrient enrichment and hydrologic status resulted in distinct bacterial communities associated with litter and bulk soils. Drier conditions and nutrient enrichment increased decomposition rates until N limitation constrains microbial community structure. This study provided the opportunity to compare the extent that community composition matters to decomposition rate – using model litter. In these nutrient poor wetland soils undergoing nutrient enrichment, the fertilization increased bacterial diversity the potential for increased C losses through decomposition is observed and is expected to increase as wetlands are drained and fertilized.

## Acknowledgments

We thank Allison Fisk, Emma Richards, Tom Vogel, John Stiller, Suelen Tullio, and Adam Gold for laboratory and field assistance. We thank John Gill and the East Carolina University grounds crew for their tireless efforts in maintaining the long-term ecological experiment. This work was supported by the East Carolina University Undergraduate Research and Creative Activity Award to MEK and the National Science Foundation (GRFP to RBB, DEB 1845845 to ALP). All code and data used in this study can be found in a public GitHub repository (https://github.com/PeraltaLab/WRC18_RhizoTeaDecomp) and the NCBI SRA (BioProject).

