## Supplemental_Material for "Distinct microbial communities alter litter decomposition rates in a fertilized coastal plain wetland"

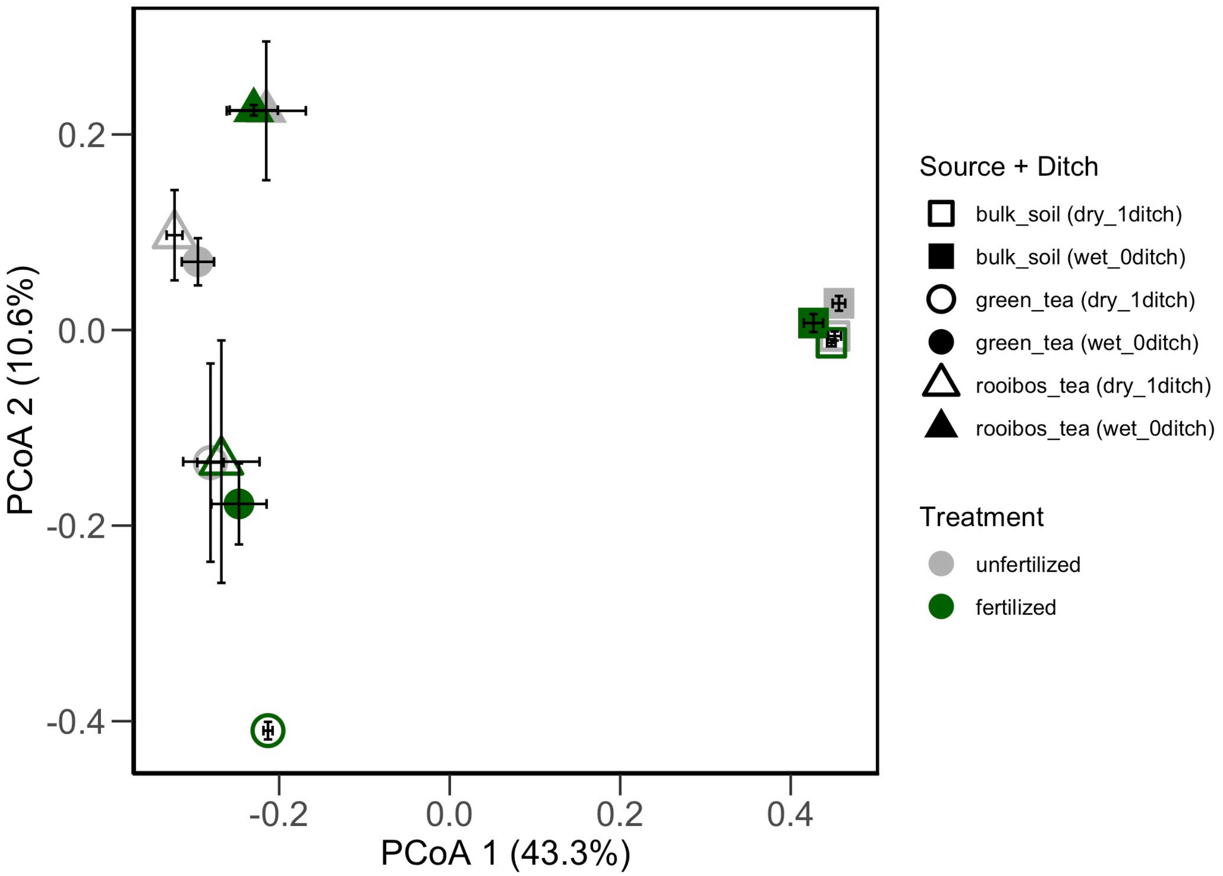


**Figure S1.** Ordination based on a Principal Coordinates Analysis depicting bacterial community composition according to tea type. Symbols are colored according to fertilization treatment (gray = unfertilized, black = fertilized) and tea source (square = bulk soil, circles = green tea, triangles = rooibos tea) at drier mowed plots situated close to the drainage ditch (open symbols) compared to wetter mowed plots (closed symbols).

**Table S1.** Summary of Type II Analysis of Variance Table with Kenward-Roger's method comparing soil properties between unfertilized and fertilized treatments. Mean (± standard deviation) of bulk soils of the interaction between fertilization treatments and ditch proximity (A). Mean (± standard deviation) of bulk soils of unfertilized (UF) and fertilized (F) treatments collected from surface soils (B). Mean (± standard deviation) of bulk soils of wetter plots away from the ditch (0Ditch) and drier plots adjacent to the (1Ditch) (C).

(A)

| Soil Factor | 0Ditch/UF | 1Ditch/UF | 0Ditch/F | 1Ditch/F | F-value | P-value |
|---|---|---|---|---|---|---|
| Temperature °C | 13.5 ± 1.3 | 11.88 ± 1.7 | 13.5 ± 1.0 | 13.0 ± 1.4 | 0.162 | 0.690 |
| Moisture (%) | 34.4 ± 0.8 | 28.6 ± 5.03 | 36.6 ± 1.1 | 30.7 ± 3.9 | 0.052 | 0.822 |
| **pH** | 3.89 ± 0.24 | 4.10 ± 0.13 | 4.06 ± 0.22 | 4.19 ± 0.06 | 4.688 | **0.037** |
| Total C (%) | 4.43 ± 0.51 | 3.67 ± 1.26 | 5.11 ± 0.89 | 4.25 ± 1.24 | 0.012 | 0.915 |
| Total N (%) | 0.23 ± 0.02 | 0.21 ± 0.06 | 0.27 ± 0.04 | 0.24 ± 0.07 | 0.012 | 0.914 |
| **Soil C:N (wt:wt)** | 19.7 ± 1.3 | 17.3 ± 0.7 | 19.1 ± 1.1 | 17.4 ± 1.1 | 7.370 | **0.010** |
| $NH_4^+$-N (µg/g dry soil) | 0.26 ± 0.26 | 0.30 ± 0.10 | 0.30 ± 0.18 | 0.26 ± 0.11 | 2.409 | 0.130 |
| **$NO_3^-$-N (µg/g dry soil)** | 0.14 ± 0.11 | MDL | 0.32 ± 0.32 | MDL | 7.913 | **0.008** |
| Total $PO_4^{3-}$-P (µg/g soil) | 123.9 ± 11.9 | 123.2 ± 31.5 | 299.4 ± 19.1 | 273.5 ± 91.8 | 0.170 | 0.694 |
| **Water ext. $PO_4^{3-}$-P (µg/g soil)** | 0.19 ± 0.16 | 0.03 ± 0.07 | 2.91 ± 1.7 | 1.04 ± 0.44 | 27.230 | **<0.001** |

(B)

| Soil Factor | Unfertilized | Fertilized | F-value | P-value |
|---|---|---|---|---|
| Temperature °C | 12.7 ± 1.6 | 13.3 ± 1.2 | 0.175 | 0.678 |
| **Moisture (%)** | 31.49 ± 4.55 | 33.66 ± 4.12 | 8.472 | **0.006** |
| **pH** | 3.99 ± 0.21 | 4.12 ± 0.16 | 49.169 | **<0.001** |
| **Total C (%)** | 4.06 ± 0.98 | 4.68 ± 1.10 | 7.501 | **0.010** |
| **Total N (%)** | 0.22 ± 0.04 | 0.26 ± 0.05 | 9.780 | **0.004** |
| Soil C:N (wt:wt) | 18.5 ± 1.6 | 18.3 ± 1.3 | 2.057 | 0.160 |
| $NH_4^+$-N (μg/g dry soil) | 0.28 ± 0.18 | 0.28 ± 0.14 | 0.116 | 0.736 |
| **$NO_3^-$-N (μg/g dry soil)** | 0.12 ± 0.10 | 0.32 ± 0.32 | 3.965 | **0.054** |
| **Total $PO_4^{3-}$-P (μg/g soil)** | 123.54 ± 22.01 | 286.45 ± 62.89 | 200.074 | **<0.001** |
| **Water ext. $PO_4^{3-}$-P (μg/g soil)** | 0.12 ± 0.14 | 1.974 ± 1.51 | 130.481 | **<0.001** |

(C)

| Soil Factor | Wet_0ditch | Dry_1ditch | F-value | P-value |
|---|---|---|---|---|
| Temperature °C | 13.5 ± 1.1 | 12.4 ± 1.5 | 2.061 | 0.201 |
| **Moisture (%)** | 35.50 ± 1.51 | 29.65 ± 4.30 | 22.810 | **0.003** |
| pH | 3.97 ± 0.23 | 4.14 ± 0.10 | 2.089 | 0.199 |
| Total C (%) | 4.77 ± 0.76 | 3.96 ± 1.20 | 4.455 | 0.080 |
| Total N (%) | 0.25 ± 0.04 | 0.25 ± 0.06 | 0.972 | 0.363 |
| **Soil C:N (wt:wt)** | 19.4 ± 1.2 | 17.4 ± 0.9 | 8.775 | **0.025** |
| $NH_4^+$-N (μg/g dry soil) | 0.28 ± 0.21 | 0.28 ± 0.10 | 0.006 | 0.942 |
| $NO_3^-$-N (μg/g dry soil) | 0.22 ± 0.22 | MDL | 3.664 | 0.104 |
| Total $PO_4^{3-}$-P (μg/g soil) | 211.6 ± 95.0 | 198.4 ± 102.4 | 0.283 | 0.605 |
| Water ext. $PO_4^{3-}$-P (μg/g soil) | 1.55 ± 1.82 | 0.53 ± 0.61 | 4.109 | 0.089 |

**Table S2.** Summary of Type II Analysis of Variance Table with Kenward-Roger's method comparing decomposition rates (A) due to source (green tea, rooibos tea), treatment (fertilized, unfertilized), and proximity to ditch (wet, dry) and soil factors (B) due to treatment and ditch.

(A)

| Fixed Effect | SumSq | MeanSq | NumDF | DenDF | F-value | Pr(>F) |
|---|---|---|---|---|---|---|
| **source** | 3489.5 | 3489.5 | 1 | 15.426 | 204.863 | **<0.001** |
| **treatment** | 380.3 | 380.3 | 1 | 15.426 | 22.329 | **<0.001** |
| ditch | 79.3 | 79.3 | 1 | 5.919 | 4.658 | 0.075 |
| **source:treatment** | 81.3 | 81.3 | 1 | 15.247 | 4.776 | **0.045** |
| source:ditch | 1.9 | 1.9 | 1 | 15.452 | 0.110 | 0.744 |
| treatment:ditch | 12.6 | 12.6 | 1 | 15.452 | 0.740 | 0.403 |
| source:treatment:ditch | 5.2 | 5.2 | 1 | 15.248 | 0.304 | 0.589 |

(B)

Temperature

| Fixed Effect | SumSq | MeanSq | NumDF | DenDF | F-value | Pr(>F) |
|---|---|---|---|---|---|---|
| treatment | 0.041 | 0.041 | 1 | 35.046 | 0.175 | 0.678 |
| ditch | 0.479 | 0.479 | 1 | 5.998 | 2.061 | 0.201 |
| treatment:ditch | 0.038 | 0.038 | 1 | 35.046 | 0.162 | 0.690 |

Moisture

| Fixed Effect | SumSq | MeanSq | NumDF | DenDF | F-value | Pr(>F) |
|---|---|---|---|---|---|---|
| treatment | 64.266 | 64.266 | 1 | 35.448 | 8.472 | **0.006** |
| ditch | 173.029 | 173.029 | 1 | 5.959 | 22.810 | **0.003** |
| treatment:ditch | 0.390 | 0.390 | 1 | 35.457 | 0.052 | 0.822 |

pH

| Fixed Effect | SumSq | MeanSq | NumDF | DenDF | F-value | Pr(>F) |
|---|---|---|---|---|---|---|
| treatment | 0.206 | 0.206 | 1 | 35.026 | 49.169 | **<0.001** |
| ditch | 0.009 | 0.009 | 1 | 5.999 | 2.089 | 0.199 |
| treatment:ditch | 0.020 | 0.020 | 1 | 35.027 | 4.688 | **0.037** |

Total C

| Fixed Effect | SumSq | MeanSq | NumDF | DenDF | F-value | Pr(>F) |
|---|---|---|---|---|---|---|
| treatment | 5.650 | 5.650 | 1 | 35.476 | 7.501 | **0.010** |
| ditch | 3.356 | 3.356 | 1 | 5.955 | 4.455 | 0.080 |
| treatment:ditch | 0.009 | 0.009 | 1 | 35.485 | 0.012 | 0.915 |

Total N

| Fixed Effect | SumSq | MeanSq | NumDF | DenDF | F-value | Pr(>F) |
|---|---|---|---|---|---|---|
| treatment | 0.01959 | 0.01959 | 1 | 35.560 | 9.780 | **0.004** |
| ditch | 0.00195 | 0.00195 | 1 | 5.942 | 0.972 | 0.363 |
| treatment:ditch | 0.00002 | 0.00002 | 1 | 35.570 | 0.012 | 0.914 |

| | Fixed Effect | SumSq | MeanSq | NumDF | DenDF | F-value | Pr(>F) |
|---|---|---|---|---|---|---|---|
| Soil C:N | treatment | 0.516 | 0.516 | 1 | 35.049 | 2.057 | 0.160 |
| | ditch | 2.202 | 2.202 | 1 | 5.997 | 8.775 | **0.025** |
| | treatment:ditch | 1.849 | 1.849 | 1 | 35.050 | 7.370 | **0.010** |

| | Fixed Effect | SumSq | MeanSq | NumDF | DenDF | F-value | Pr(>F) |
|---|---|---|---|---|---|---|---|
| Ammonium | treatment | 0.0004 | 0.0004 | 1 | 35.024 | 0.116 | 0.736 |
| | ditch | 0.0000 | 0.0000 | 1 | 5.999 | 0.006 | 0.942 |
| | treatment:ditch | 0.0092 | 0.0092 | 1 | 35.025 | 2.409 | 0.130 |

| | Fixed Effect | SumSq | MeanSq | NumDF | DenDF | F-value | Pr(>F) |
|---|---|---|---|---|---|---|---|
| Nitrate | treatment | 0.030 | 0.030 | 1 | 35.070 | 3.965 | **0.054** |
| | ditch | 0.028 | 0.028 | 1 | 5.996 | 3.664 | 0.104 |
| | treatment:ditch | 0.060 | 0.060 | 1 | 35.072 | 7.913 | **0.008** |

| | Fixed Effect | SumSq | MeanSq | NumDF | DenDF | F-value | Pr(>F) |
|---|---|---|---|---|---|---|---|
| Total $PO_4^{3-}$-P | treatment | 304321 | 304321 | 1 | 35.294 | 200.074 | **<0.001** |
| | ditch | 259 | 259 | 1 | 5.978 | 0.170 | 0.694 |
| | treatment:ditch | 894 | 894 | 1 | 35.299 | 0.588 | 0.448 |

| | Fixed Effect | SumSq | MeanSq | NumDF | DenDF | F-value | Pr(>F) |
|---|---|---|---|---|---|---|---|
| Water ext. $PO_4^{3-}$-P | treatment | 39.505 | 39.505 | 1 | 35.113 | 130.481 | **<0.001** |
| | ditch | 1.244 | 1.244 | 1 | 5.993 | 4.109 | 0.089 |
| | treatment:ditch | 8.244 | 8.244 | 1 | 35.115 | 27.230 | **<0.001** |

**Table S3.** Summary PERMANOVA comparing bacterial communities due to source/tea type (green tea, rooibos tea), treatment (unfertilized, fertilized), and proximity to ditch (wet, dry).

| Factor | df | SumsOfSqs | MeanSqs | F.Model | $R^2$ | Pr(>F) |
|---|---|---|---|---|---|---|
| **source** | 1 | 0.710 | 0.710 | 4.361 | 0.125 | **0.001** |
| **treatment** | 1 | 0.377 | 0.377 | 2.318 | 0.067 | **0.006** |
| **ditch** | 1 | 0.594 | 0.594 | 3.647 | 0.105 | **0.001** |
| source:treatment | 1 | 0.198 | 0.198 | 1.216 | 0.035 | 0.190 |
| **source:ditch** | 1 | 0.340 | 0.340 | 2.086 | 0.060 | **0.013** |
| treatment:ditch | 1 | 0.197 | 0.197 | 1.211 | 0.035 | 0.204 |
| source:treatment:ditch | 1 | 0.162 | 0.162 | 0.992 | 0.028 | 0.396 |
| Residuals | 19 | 3.093 | 0.163 | 0.545 | | |
| Total | 26 | 5.670 | 1.000 | | | |

**Table S4.** Summary PERMANOVA comparing bacterial communities due to source (bulk soil, green tea, rooibos tea), treatment (unfertilized, fertilized), and proximity to ditch (wet, dry).

| Factor | df | SumsOfSqs | MeanSqs | F.Model | $R^2$ | Pr(>F) |
|---|---|---|---|---|---|---|
| **source** | 2 | 5.776 | 2.888 | 23.947 | 0.486 | **0.001** |
| **treatment** | 1 | 0.371 | 0.370 | 3.072 | 0.031 | **0.019** |
| **ditch** | 1 | 0.471 | 0.471 | 3.905 | 0.040 | **0.008** |
| **source:treatment** | 2 | 0.469 | 0.235 | 1.945 | 0.039 | **0.037** |
| **source:ditch** | 2 | 0.640 | 0.320 | 2.652 | 0.054 | **0.004** |
| treatment:ditch | 1 | 0.151 | 0.151 | 1.254 | 0.013 | 0.228 |
| source:treatment:ditch | 2 | 0.268 | 0.134 | 1.111 | 0.023 | 0.291 |
| Residuals | 31 | 3.739 | 0.121 | 0.315 | | |
| Total | 42 | 11.884 | 1.000 | | | |

**Table S5.** Summary of Type II Analysis of Variance Table with Kenward-Roger's method comparing bacterial diversity metrics (OTU richness, Shannon Diversity Index H′, and Simpson's evenness) associated with source (bulk soil, green tea, rooibos tea), treatment (unfertilized, fertilized), and proximity to ditch (wet, dry).

(A) OTU Richness

| Fixed Effect | SumSq | MeanSq | NumDF | DenDF | F-value | Pr(>F) |
|---|---|---|---|---|---|---|
| **source** | 27269829 | 13634915 | 2 | 25.585 | 92.599 | **<0.0001** |
| **treatment** | 1189815 | 1189815 | 1 | 25.469 | 8.081 | **0.009** |
| ditch | 47449 | 47449 | 1 | 5.823 | 0.322 | 0.591 |
| source:treatment | 202762 | 101381 | 2 | 25.720 | 0.689 | 0.511 |
| source:ditch | 302781 | 151391 | 2 | 25.687 | 1.028 | 0.372 |
| treatment:ditch | 41789 | 41789 | 1 | 25.465 | 0.284 | 0.599 |

(B) Shannon Diversity

| Fixed Effect | SumSq | MeanSq | NumDF | DenDF | F-value | Pr(>F) |
|---|---|---|---|---|---|---|
| **source** | 8.554 | 4.277 | 2 | 25.760 | 35.117 | **<0.0001** |
| **treatment** | 2.349 | 2.349 | 1 | 25.621 | 19.289 | **<0.0001** |
| ditch | 0.553 | 0.553 | 1 | 5.762 | 4.539 | 0.079 |
| source:treatment | 0.556 | 0.278 | 2 | 25.936 | 2.284 | 0.122 |
| source:ditch | 0.395 | 0.198 | 2 | 25.889 | 1.623 | 0.217 |
| treatment:ditch | 0.035 | 0.035 | 1 | 25.616 | 0.287 | 0.597 |
| source:treatment:ditch | 0.161 | 0.080 | 2 | 26.007 | 0.660 | 0.526 |

(C) Simpson's Evenness

| Fixed Effect | SumSq | MeanSq | NumDF | DenDF | F-value | Pr(>F) |
|---|---|---|---|---|---|---|
| **source** | 0.0023 | 0.0011 | 2 | 25.698 | 3.929 | **0.032** |
| **treatment** | 0.0026 | 0.0026 | 1 | 25.566 | 8.958 | **0.006** |
| ditch | 0.0006 | 0.0006 | 1 | 5.784 | 2.124 | 0.197 |
| source:treatment | 0.0007 | 0.0003 | 2 | 25.859 | 1.184 | 0.322 |
| source:ditch | 0.0011 | 0.0005 | 2 | 25.817 | 1.885 | 0.172 |
| treatment:ditch | 0.0001 | 0.0001 | 1 | 25.561 | 0.394 | 0.536 |
| source:treatment:ditch | 0.0003 | 0.0002 | 2 | 25.925 | 0.552 | 0.582 |

**Table S6.** Bacterial taxa (OTUs) representing the unique taxa associated to treatment type according to indicator species analysis. This summary represents the top bacterial taxa associated with each treatment type.

Bulk soil

| | dry_1ditch | wet_0ditch |
|---|---|---|
| unfertilized | Proteobacteria/Alphaproteobacteria/Rhodospirillales/Rhodospirillales_unclassified/Rhodospirillales_unclassified | Verrucomicrobia/Spartobacteria/Spartobacteria_unclassified/Spartobacteria_unclassified/Spartobacteria_unclassified<br>Acidobacteria/Acidobacteria_Gp1/Gp1/Gp1_unclassified/Gp1_unclassified |
| fertilized | Acidobacteria/Acidobacteria_Gp2/Gp2/Gp2_unclassified/Gp2_unclassified<br>…Gp1_unclassified<br>…Gp6_unclassified | Actinobacteria/Actinobacteria/Solirubrobacterales/Solirubrobacterales_unclassified/Solirubrobacterales_unclassified |

Green tea

| | dry_1ditch | wet_0ditch |
|---|---|---|
| unfertilized | Proteobacteria/Alphaproteobacteria/Caulobacterales/Caulobacteraceae/Phenylobacterium | Acidobacteria/Acidobacteria_Gp3/Gp3/Gp3_unclassified/Gp3_unclassified<br>Proteobacteria/Alphaproteobacteria/Rhodospirillales/Rhodospirillaceae/Dongia |
| fertilized | Proteobacteria/Gammaproteobacteria/Legionellales/Legionellaceae/Legionella | Actinobacteria/Actinobacteria/Solirubrobacterales/Conexibacteraceae/Conexibacter |

Rooibos tea

| | dry_1ditch | wet_0ditch |
|---|---|---|
| unfertilized | Proteobacteria/Alphaproteobacteria/Rhodospirillales/Rhodospirillaceae/Lacibacterium | Proteobacteria/Alphaproteobacteria/Rhodospirillales/Acetobacteraceae/Acidisoma |
| fertilized | Proteobacteria/Gammaproteobacteria/Xanthomonadales/Xanthomonadaceae/Dokdonella<br>Actinobacteria/Actinobacteria/Actinomycetales/Microbacteriaceae/Microbacteriaceae_unclassified | Proteobacteria/Gammaproteobacteria/Xanthomonadales/Xanthomonadaceae/Dyella |

**Table S7a.** Bacterial taxa (OTUs) with the highest individual value representing the unique taxa associated to treatment type according to indicator species analysis. This summary represents the top bacterial taxa associated with each treatment type (a) and all the taxa that are significantly representative of the treatment types (b).

| OTU | Cluster | Cluster_ID | IndVal | Prob | Phylum/Class/Order/Family/Genus |
|---|---|---|---|---|---|
| Otu00006 | 1 | bulk_soil.M.0 | 0.443 | 0.001 | Verrucomicrobia/Spartobacteria/Spartobacteria_unclassified/Spartobacteria_unclassified/Spartobacteria_unclassified |
| Otu00056 | 1 | bulk_soil.M.0 | 0.432 | 0.014 | Acidobacteria/Acidobacteria_Gp1/Gp1/Gp1_unclassified/Gp1_unclassified |
| Otu00181 | 2 | green_tea.M.0 | 0.507 | 0.003 | Acidobacteria/Acidobacteria_Gp3/Gp3/Gp3_unclassified/Gp3_unclassified |
| Otu00076 | 2 | green_tea.M.0 | 0.426 | 0.012 | Proteobacteria/Alphaproteobacteria/Rhodospirillales/Rhodospirillaceae/Dongia |
| Otu00092 | 3 | rooibos_tea.M.0 | 0.625 | 0.014 | Proteobacteria/Alphaproteobacteria/Rhodospirillales/Acetobacteraceae/Acidisoma |
| Otu00050 | 4 | bulk_soil.MF.0 | 0.403 | 0.001 | Actinobacteria/Actinobacteria/Solirubrobacterales/Solirubrobacterales_unclassified/Solirubrobacterales_unclassified |
| Otu00127 | 5 | green_tea.MF.0 | 0.501 | 0.046 | Actinobacteria/Actinobacteria/Solirubrobacterales/Conexibacteraceae/Conexibacter |
| Otu00031 | 6 | rooibos_tea.MF.0 | 0.347 | 0.028 | Proteobacteria/Gammaproteobacteria/Xanthomonadales/Xanthomonadaceae/Dyella |
| Otu00081 | 7 | bulk_soil.M.1 | 0.437 | 0.001 | Proteobacteria/Alphaproteobacteria/Rhodospirillales/Rhodospirillales_unclassified/Rhodospirillales_unclassified |
| Otu00185 | 8 | green_tea.M.1 | 0.401 | 0.032 | Proteobacteria/Alphaproteobacteria/Caulobacterales/Caulobacteraceae/Phenylobacterium |
| Otu00173 | 9 | rooibos_tea.M.1 | 0.699 | 0.007 | Proteobacteria/Alphaproteobacteria/Rhodospirillales/Rhodospirillaceae/Lacibacterium |
| Otu00078 | 10 | bulk_soil.MF.1 | 0.376 | 0.003 | Acidobacteria/Acidobacteria_Gp2/Gp2/Gp2_unclassified/Gp2_unclassified |
| Otu00036 | 10 | bulk_soil.MF.1 | 0.347 | 0.001 | Acidobacteria/Acidobacteria_Gp1/Gp1/Gp1_unclassified/Gp1_unclassified |
| Otu00045 | 10 | bulk_soil.MF.1 | 0.335 | 0.005 | Acidobacteria/Acidobacteria_Gp6/Gp6/Gp6_unclassified/Gp6_unclassified |
| Otu00075 | 11 | green_tea.MF.1 | 0.716 | 0.007 | Proteobacteria/Gammaproteobacteria/Legionellales/Legionellaceae/Legionella |
| Otu00108 | 12 | rooibos_tea.MF.1 | 0.347 | 0.048 | Proteobacteria/Gammaproteobacteria/Xanthomonadales/Xanthomonadaceae/Dokdonella |
| Otu00166 | 12 | rooibos_tea.MF.1 | 0.337 | 0.016 | Actinobacteria/Actinobacteria/Actinomycetales/Microbacteriaceae/Microbacteriaceae_unclassified |

**Table S7b.** Complete list of bacterial taxa (OTUs) unique to treatment type according to indicator species analysis. This summary represents the top bacterial taxa associated with each source (bulk soil, green tea, rooibos tea), treatment (M: unfertilized; MF: fertilized), and proximity to ditch (0: wet, far from ditch; 1: dry, close to ditch). Rows highlighted in light green are the same rows presented in Table S7a.

| OTU | Cluster | Cluster_ID | IndVal | Prob | Phylum/Class/Order/Family/Genus |
|---|---|---|---|---|---|
| Otu00006 | 1 | bulk_soil.M.0 | 0.443 | 0.001 | Verrucomicrobia/Spartobacteria/Spartobacteria_unclassified/Spartobacteria_unclassified/Spartobacteria_unclassified |
| Otu00056 | 1 | bulk_soil.M.0 | 0.432 | 0.014 | Acidobacteria/Acidobacteria_Gp1/Gp1/Gp1_unclassified/Gp1_unclassified |
| Otu00030 | 1 | bulk_soil.M.0 | 0.412 | 0.002 | Planctomycetes/Planctomycetia/Planctomycetales/Planctomycetaceae/Planctomycetaceae_unclassified |
| Otu00048 | 1 | bulk_soil.M.0 | 0.401 | 0.001 | Proteobacteria/Alphaproteobacteria/Rhizobiales/Hyphomicrobiaceae/Rhodomicrobium |
| Otu00035 | 1 | bulk_soil.M.0 | 0.375 | 0.001 | Acidobacteria/Acidobacteria_Gp1/Gp1/Gp1_unclassified/Gp1_unclassified |
| Otu00025 | 1 | bulk_soil.M.0 | 0.329 | 0.015 | Verrucomicrobia/Spartobacteria/Spartobacteria_unclassified/Spartobacteria_unclassified/Spartobacteria_unclassified |
| Otu00053 | 1 | bulk_soil.M.0 | 0.323 | 0.011 | Acidobacteria/Acidobacteria_Gp13/Gp13/Gp13_unclassified/Gp13_unclassified |
| Otu00049 | 1 | bulk_soil.M.0 | 0.304 | 0.021 | Acidobacteria/Acidobacteria_Gp1/Gp1/Gp1_unclassified/Gp1_unclassified |
| Otu00008 | 1 | bulk_soil.M.0 | 0.303 | 0.026 | Verrucomicrobia/Spartobacteria/Spartobacteria_unclassified/Spartobacteria_unclassified/Spartobacteria_unclassified |
| Otu00047 | 1 | bulk_soil.M.0 | 0.286 | 0.014 | Bacteria_unclassified/Bacteria_unclassified/Bacteria_unclassified/Bacteria_unclassified/Bacteria_unclassified |
| Otu00005 | 1 | bulk_soil.M.0 | 0.285 | 0.002 | Proteobacteria/Alphaproteobacteria/Rhizobiales/Roseiarcaceae/Roseiarcus |
| Otu00062 | 1 | bulk_soil.M.0 | 0.278 | 0.008 | Acidobacteria/Acidobacteria_Gp1/Gp1/Gp1_unclassified/Gp1_unclassified |
| Otu00013 | 1 | bulk_soil.M.0 | 0.278 | 0.027 | Acidobacteria/Acidobacteria_Gp2/Gp2/Gp2_unclassified/Gp2_unclassified |
| Otu00028 | 1 | bulk_soil.M.0 | 0.276 | 0.016 | Acidobacteria/Acidobacteria_Gp2/Gp2/Gp2_unclassified/Gp2_unclassified |
| Otu00181 | 2 | green_tea.M.0 | 0.507 | 0.003 | Acidobacteria/Acidobacteria_Gp3/Gp3/Gp3_unclassified/Gp3_unclassified |
| Otu00076 | 2 | green_tea.M.0 | 0.426 | 0.012 | Proteobacteria/Alphaproteobacteria/Rhodospirillales/Rhodospirillaceae/Dongia |
| Otu00197 | 2 | green_tea.M.0 | 0.360 | 0.046 | Proteobacteria/Alphaproteobacteria/Rhodospirillales/Rhodospirillaceae/Rhodospirillaceae_unclassified |

| | | | | | |
|---|---|---|---|---|---|
| Otu00029 | 2 | green_tea.M.0 | 0.357 | 0.024 | Proteobacteria/Alphaproteobacteria/Rhodospirillales/Rhodospirillaceae/Rhodospirillaceae_unclassified |
| Otu00003 | 2 | green_tea.M.0 | 0.296 | 0.042 | Proteobacteria/Betaproteobacteria/Burkholderiales/Burkholderiaceae/Burkholderia |
| Otu00093 | 2 | green_tea.M.0 | 0.288 | 0.048 | Bacteroidetes/Sphingobacteriia/Sphingobacteriales/Chitinophagaceae/Chitinophagaceae_unclassified |
| Otu00041 | 2 | green_tea.M.0 | 0.284 | 0.024 | Verrucomicrobia/Spartobacteria/Spartobacteria_unclassified/Spartobacteria_unclassified/Spartobacteria_unclassified |
| Otu00019 | 2 | green_tea.M.0 | 0.257 | 0.014 | Verrucomicrobia/Subdivision3/Subdivision3_unclassified/Subdivision3_unclassified/Subdivision3_unclassified |
| Otu00092 | 3 | rooibos_tea.M.0 | 0.625 | 0.014 | Proteobacteria/Alphaproteobacteria/Rhodospirillales/Acetobacteraceae/Acidisoma |
| Otu00064 | 3 | rooibos_tea.M.0 | 0.465 | 0.033 | Proteobacteria/Alphaproteobacteria/Rhodospirillales/Rhodospirillaceae/Rhodospirillaceae_unclassified |
| Otu00077 | 3 | rooibos_tea.M.0 | 0.343 | 0.001 | Proteobacteria/Alphaproteobacteria/Caulobacterales/Caulobacteraceae/Phenylobacterium |
| Otu00040 | 3 | rooibos_tea.M.0 | 0.340 | 0.027 | Proteobacteria/Gammaproteobacteria/Gammaproteobacteria_unclassified/Gammaproteobacteria_unclassified/Gammaproteobacteria_unclassified |
| Otu00050 | 4 | bulk_soil.MF.0 | 0.403 | 0.001 | Actinobacteria/Actinobacteria/Solirubrobacterales/Solirubrobacterales_unclassified/Solirubrobacterales_unclassified |
| Otu00020 | 4 | bulk_soil.MF.0 | 0.362 | 0.001 | Actinobacteria/Actinobacteria/Solirubrobacterales/Solirubrobacterales_unclassified/Solirubrobacterales_unclassified |
| Otu00073 | 4 | bulk_soil.MF.0 | 0.342 | 0.022 | Acidobacteria/Acidobacteria_Gp2/Gp2/Gp2_unclassified/Gp2_unclassified |
| Otu00018 | 4 | bulk_soil.MF.0 | 0.335 | 0.001 | Actinobacteria/Actinobacteria/Actinomycetales/Thermomonosporaceae/Actinoallomurus |
| Otu00085 | 4 | bulk_soil.MF.0 | 0.312 | 0.024 | Planctomycetes/Planctomycetia/Planctomycetales/Planctomycetaceae/Planctomycetaceae_unclassified |
| Otu00007 | 4 | bulk_soil.MF.0 | 0.283 | 0.005 | Acidobacteria/Acidobacteria_Gp1/Gp1/Gp1_unclassified/Gp1_unclassified |
| Otu00060 | 4 | bulk_soil.MF.0 | 0.256 | 0.006 | Actinobacteria/Actinobacteria/Actinomycetales/Actinomycetales_unclassified/Actinomycetales_unclassified |
| Otu00044 | 4 | bulk_soil.MF.0 | 0.234 | 0.019 | Verrucomicrobia/Subdivision3/Subdivision3_unclassified/Subdivision3_unclassified/Subdivision3_unclassified |
| Otu00127 | 5 | green_tea.MF.0 | 0.501 | 0.046 | Actinobacteria/Actinobacteria/Solirubrobacterales/Conexibacteraceae/Conexibacter |
| Otu00065 | 5 | green_tea.MF.0 | 0.320 | 0.049 | Proteobacteria/Betaproteobacteria/Burkholderiales/Comamonadaceae/Comamonadaceae_unclassified |
| Otu00031 | 6 | rooibos_tea.MF.0 | 0.347 | 0.028 | Proteobacteria/Gammaproteobacteria/Xanthomonadales/Xanthomonadaceae/Dyella |

| | | | | | |
|---|---|---|---|---|---|
| Otu00115 | 6 | rooibos_tea.MF.0 | 0.314 | 0.028 | Verrucomicrobia/Subdivision3/Subdivision3_unclassified/Subdivision3_unclassified/Subdivision3_unclassified |
| Otu00109 | 6 | rooibos_tea.MF.0 | 0.304 | 0.042 | Proteobacteria/Gammaproteobacteria/Gammaproteobacteria_unclassified/Gammaproteobacteria_unclassified/Gammaproteobacteria_unclassified |
| Otu00138 | 6 | rooibos_tea.MF.0 | 0.257 | 0.028 | Proteobacteria/Alphaproteobacteria/Caulobacterales/Caulobacteraceae/Phenylobacterium |
| Otu00066 | 6 | rooibos_tea.MF.0 | 0.234 | 0.046 | Proteobacteria/Alphaproteobacteria/Rhodospirillales/Acetobacteraceae/Acetobacteraceae_unclassified |
| Otu00081 | 7 | bulk_soil.M.1 | 0.437 | 0.001 | Proteobacteria/Alphaproteobacteria/Rhodospirillales/Rhodospirillales_unclassified/Rhodospirillales_unclassified |
| Otu00014 | 7 | bulk_soil.M.1 | 0.351 | 0.001 | Proteobacteria/Alphaproteobacteria/Rhizobiales/Rhizobiales_unclassified/Rhizobiales_unclassified |
| Otu00100 | 7 | bulk_soil.M.1 | 0.347 | 0.001 | Proteobacteria/Alphaproteobacteria/Rhodospirillales/Rhodospirillales_unclassified/Rhodospirillales_unclassified |
| Otu00009 | 7 | bulk_soil.M.1 | 0.346 | 0.001 | Verrucomicrobia/Spartobacteria/Spartobacteria_unclassified/Spartobacteria_unclassified/Spartobacteria_unclassified |
| Otu00043 | 7 | bulk_soil.M.1 | 0.323 | 0.018 | Acidobacteria/Acidobacteria_Gp1/Gp1/Gp1_unclassified/Gp1_unclassified |
| Otu00089 | 7 | bulk_soil.M.1 | 0.287 | 0.018 | Proteobacteria/Proteobacteria_unclassified/Proteobacteria_unclassified/Proteobacteria_unclassified/Proteobacteria_unclassified |
| Otu00032 | 7 | bulk_soil.M.1 | 0.266 | 0.007 | Actinobacteria/Actinobacteria/Solirubrobacterales/Conexibacteraceae/Conexibacter |
| Otu00017 | 7 | bulk_soil.M.1 | 0.264 | 0.019 | Acidobacteria/Acidobacteria_Gp2/Gp2/Gp2_unclassified/Gp2_unclassified |
| Otu00058 | 7 | bulk_soil.M.1 | 0.264 | 0.008 | Verrucomicrobia/Subdivision3/Subdivision3_unclassified/Subdivision3_unclassified/Subdivision3_unclassified |
| Otu00004 | 7 | bulk_soil.M.1 | 0.252 | 0.002 | Proteobacteria/Alphaproteobacteria/Rhizobiales/Rhizobiales_unclassified/Rhizobiales_unclassified |
| Otu00051 | 7 | bulk_soil.M.1 | 0.247 | 0.010 | Actinobacteria/Actinobacteria/Actinomycetales/Mycobacteriaceae/Mycobacterium |
| Otu00185 | 8 | green_tea.M.1 | 0.401 | 0.032 | Proteobacteria/Alphaproteobacteria/Caulobacterales/Caulobacteraceae/Phenylobacterium |
| Otu00173 | 9 | rooibos_tea.M.1 | 0.699 | 0.007 | Proteobacteria/Alphaproteobacteria/Rhodospirillales/Rhodospirillaceae/Lacibacterium |
| Otu00159 | 9 | rooibos_tea.M.1 | 0.311 | 0.012 | Proteobacteria/Alphaproteobacteria/Rhizobiales/Rhizobiales_unclassified/Rhizobiales_unclassified |
| Otu00052 | 9 | rooibos_tea.M.1 | 0.260 | 0.021 | Proteobacteria/Alphaproteobacteria/Rhodospirillales/Rhodospirillales_unclassified/Rhodospirillales_unclassified |
| Otu00078 | 10 | bulk_soil.MF.1 | 0.376 | 0.003 | Acidobacteria/Acidobacteria_Gp2/Gp2/Gp2_unclassified/Gp2_unclassified |
| Otu00036 | 10 | bulk_soil.MF.1 | 0.347 | 0.001 | Acidobacteria/Acidobacteria_Gp1/Gp1/Gp1_unclassified/Gp1_unclassified |

| | | | | | |
|---|---|---|---|---|---|
| Otu00045 | 10 | bulk_soil.MF.1 | 0.335 | 0.005 | Acidobacteria/Acidobacteria_Gp6/Gp6/Gp6_unclassified/Gp6_unclassified |
| Otu00057 | 10 | bulk_soil.MF.1 | 0.263 | 0.022 | Acidobacteria/Acidobacteria_Gp3/Gp3/Gp3_unclassified/Gp3_unclassified |
| Otu00075 | 11 | green_tea.MF.1 | 0.716 | 0.007 | Proteobacteria/Gammaproteobacteria/Legionellales/Legionellaceae/Legionella |
| Otu00223 | 11 | green_tea.MF.1 | 0.486 | 0.014 | Proteobacteria/Alphaproteobacteria/Rhodospirillales/Rhodospirillaceae/Dongia |
| Otu00191 | 11 | green_tea.MF.1 | 0.411 | 0.002 | Proteobacteria/Alphaproteobacteria/Rhizobiales/Rhizobiales_unclassified/Rhizobiales_unclassified |
| Otu00144 | 11 | green_tea.MF.1 | 0.356 | 0.022 | Proteobacteria/Betaproteobacteria/Burkholderiales/Comamonadaceae/Pelomonas |
| Otu00152 | 11 | green_tea.MF.1 | 0.320 | 0.037 | Proteobacteria/Alphaproteobacteria/Rhizobiales/Hyphomicrobiaceae/Devosia |
| Otu00108 | 12 | rooibos_tea.MF.1 | 0.347 | 0.048 | Proteobacteria/Gammaproteobacteria/Xanthomonadales/Xanthomonadaceae/Dokdonella |
| Otu00166 | 12 | rooibos_tea.MF.1 | 0.337 | 0.016 | Actinobacteria/Actinobacteria/Actinomycetales/Microbacteriaceae/Microbacteriaceae_unclassified |
| Otu00129 | 12 | rooibos_tea.MF.1 | 0.302 | 0.033 | Bacteria_unclassified/Bacteria_unclassified/Bacteria_unclassified/Bacteria_unclassified/Bacteria_unclassified |
| Otu00090 | 12 | rooibos_tea.MF.1 | 0.296 | 0.020 | Proteobacteria/Deltaproteobacteria/Myxococcales/Myxococcales_unclassified/Myxococcales_unclassified |
| Otu00136 | 12 | rooibos_tea.MF.1 | 0.265 | 0.020 | Proteobacteria/Alphaproteobacteria/Alphaproteobacteria_unclassified/Alphaproteobacteria_unclassified/Alphaproteobacteria_unclassified |
